# Modelling the spread and mitigation of an emerging vector-borne pathogen; citrus greening in the U.S.

**DOI:** 10.1101/2022.05.04.490566

**Authors:** Viet-Anh Nguyen, David W. Bartels, Christopher A. Gilligan

## Abstract

Predictive models, based upon epidemiological principles and fitted to surveillance data, play an increasingly important role in shaping regulatory and operational policies for emerging outbreaks. Data for parameterising these strategically important models are often scarce when rapid actions are required to change the course of an epidemic invading a new region. We provide a flexible toolkit for landscape-scale disease management, which is applicable to a range of emerging pathogens including vector-borne pathogens for both endemic and invading epidemic vectors. We use the toolkit to analyse and predict the spread of Huanglongbing disease or citrus greening in the U.S. We estimate epidemiological parameters using survey data from one region (Texas) and show how to transfer and test parameters to construct predictive spatio-temporal models for another region (California). The models are used to screen effective coordinated and reactive management strategies for different regions.

## INTRODUCTION

A rapid response to emerging epidemics of crop disease is essential for successful management of an epidemic (Gilligan, 2007; Gilligan & Bosch, 2008; Meentemeyer, et al., 2011) just as for the current pandemic of COVID-19 (Wu, et al., 2020; Zhuang, et al., 2020). Effective management, however, requires knowledge of critical epidemiological parameters for transmission and dispersal of the pathogen (Adrakey, et al., 2017; Neri, et al., 2014; Parry, et al., 2014). The epidemiological parameters are required for models to predict the current extent of infection, the likely future spread (Cunniffe, et al., 2015; Filipe, et al., 2012; Neri, et al., 2014) and the effectiveness of potential intervention strategies (Cunniffe, et al., 2016; Neri, et al., 2014; Taylor, et al., 2016; Kissler, et al., 2020). The parameters are, however, seldom known when an epidemic invades a new region. There are two options to obtain parameters: wait until there are adequate surveillance data in the newly-invaded region from which to estimate parameters or transfer parameters derived for another region and incorporate them into models that allow for different host distributions and environmental variables in the newly-invaded region. Here we introduce and test a framework for an emerging epidemic of one of the most serious threats to citrus production world-wide. We adapt a model parameterised for one region and apply the model to test intervention strategies in a markedly different region. Our approach has wide application to a broad range of crop pests and pathogens that threaten food security and natural ecosystems.

Huanglongbing (HLB) disease causes severe chlorosis of foliage, dieback, loss of yield, discolouration and ill-flavour of fruit, and death of citrus trees (Gottwald, 2010). The disease is associated with three bacterial strains of which *Candidatus* Liberibacter asiaticus (*C*Las) is the prevalent type in the Western Hemisphere. The pathogen is transmitted by insect vectors and by movement of infected planting material. Options for control include quarantine, pesticide application to kill the vector, and removal of infected and surrounding trees. Quarantine involves restricting movement of planting material and citrus fruit around infected sites with radii of up to several kilometres.

HLB has caused a 74% drop in citrus production in Florida since HLB first detection in 2005 (United States Department of Agriculture, National Agricultural Statistics Service [USDA/NASS], n.d.). It has spread rapidly in Texas and has been introduced and become established in Southern California. There is a consequent risk of invasion into the major citrus production area with the Central Valley in California (Warnert, 2012), where the insect vector, the Asian citrus psyllid (ACP), *Diaphorina citri* Kuwayama, is currently invading. The rapid spread of the disease in Florida, following the introduction of the ACP vector, poses a serious threat to citrus production in the Central Valley. The pathogen also continues to pose a major threat to citrus production in Brazil (Filho, et al., 2016). The invasion of another vector, the African citrus psyllid, *Trioza erytreae* (Del Guercio), in Portugal and Spain constitutes a threat of introduction of the disease into European countries (Cocuzza, et al., 2016). The disease is also widespread in South East Asia (Bové, 2006) and the Las bacterium has recently been reported in Kenya (Ajene, et al., 2020).

Given the widespread distribution and continued spread of the pathogen and its vectors there is an urgent need for a flexible parameterised model that can be used to predict spread at landscape scales and, to inform surveillance and management options. Here we focus on the threat to citrus production in California for which there are emerging surveillance data for the pathogen and the vector. Specifically, we show how a flexible epidemic model parameterised and tested using surveillance data in one region (Texas) can be adapted to predict spread and assess options for management in a new region with different climatic conditions, where the vector is endemic (Southern California) or invading (Central Valley, California). The parameterisation allows for multiple sources for introduction of infection to a region, the integration of pathogen and vector dynamics, heterogeneous distribution of the citrus host, encompassing plantations and backyard trees. The approach also allows for incomplete surveys as well as the confounding effects on pathogen spread of pesticide application by some growers to manage the vector. We use the model to analyse the effectiveness of ACP control strategies retrospectively for the epidemic in Texas and prospectively to delay the emerging epidemic in California, as well as comparing different quarantine scenarios. The approach is readily adaptable for other crop pathosystems.

## RESULTS

### Modelling landscape-scale HLB spread in Texas and Bayesian estimation of parameters from statewide survey data

Although intensive surveillance data are available in Florida, the speed of the epidemic was such that maps of disease spread reflect more closely the timing of surveys rather than the underlying dynamics of the epidemic. Accordingly, those data were not considered suitable for parameter estimation and we used instead data from more structured surveys in Texas. We developed a continuous-time, spatially-explicit, stochastic epidemiological model (Fig. 2A) of HLB at the landscape scale, and estimated model parameters from HLB survey data. The model operates on a 1-km^2^ resolution gridded citrus landscape of the Lower Rio Grande Valley, covering commercial orchards and residential trees in 19 cities of four counties in the southernmost tip of South Texas (Fig. 1A). We assume that the exposure of healthy citrus trees to HLB infection occurs via three sources: secondary transmission effected by ACP vector movements within the Texas landscape, and two sources of primary transmission. One involves the introduction of infected trees by trade or other human-mediated movements. The other source of primary infection involves the arrival of infected ACP vectors from outside the region, including across the border with Mexico. The rates of infection are assumed to be inversely proportional to the intensity of control measures deployed to a local area. Throughout the outbreak in Texas, commercial growers voluntarily participated in an annual coordinated insecticide spray program aiming to reduce ACP densities in citrus groves, while residential trees were left mainly untreated. Survey data between December 2011 and October 2018 were utilised and comprise the location and time of collection and qPCR diagnostic results of leaf sub-samples (Fig. 1D). Although the times at which cells get exposed and infectious were not observable, we developed a data-augmented Markov chain Monte Carlo (DA-MCMC) algorithm (Gibson & Renshaw, 1998; Pooley, et al., 2015) within a Bayesian inference framework to approximate the joint posterior distributions of parameters. We designated the data from December 2011 to August 2016 as the training data set and used samples collected between September 2016 and October 2018 as a testing data set for model validation. Since the testing data were not used for model fitting, they provide an independent source to verify the capability of the model to make inferences into the future.

**Fig. 1.**
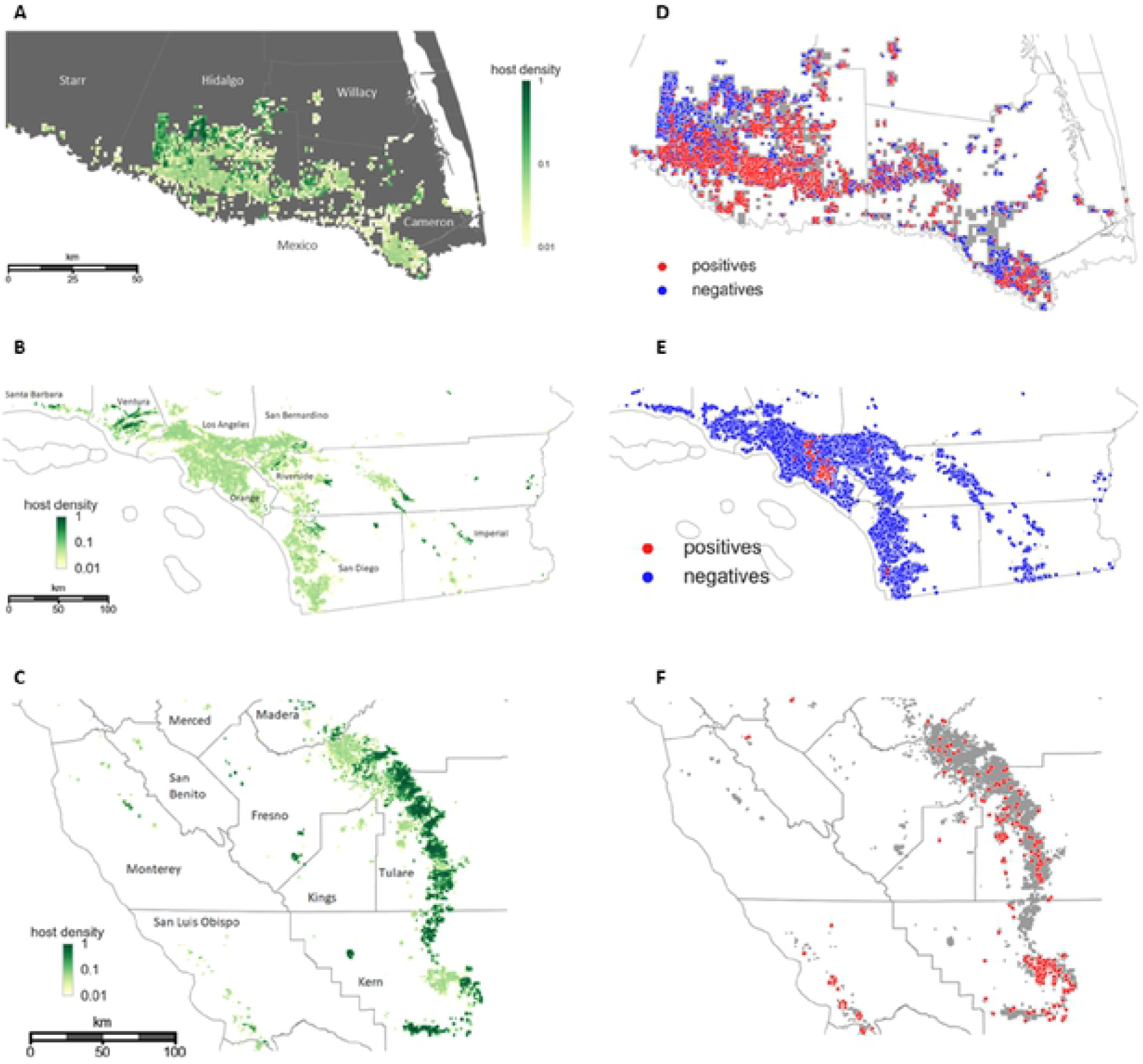
The citrus landscapes for three regions in Texas and California and the survey data for emerging HLB and ACP epidemics in the regions. **(A-C)** Gridded approximation of the citrus distribution in the (A) Lower Rio Grande Valley, Texas, (B) southern California, and (C) the Central Valley. The regions comprise large commercial citrus groves (high host density) interspersed with dooryard trees in residential areas (low host density). **(D-E)** Geo-coded diagnostic plant samples collected as part of the (D) Texas HLB state-wide survey between December 2011 and October 2018, (E) California state-wide survey between June 2015 and June 2019. We classed samples with Ct value less than 36 (out of 40 qPCR cycles) as positives. **(F)** Geo-coded ACP samples recorded by California Department of Food and Agriculture independently from the state-wide HLB survey. The samples were found in sticky yellow traps set up near trees in the Central Valley from 2012 to 2017 inclusively.

**Fig. 2.**
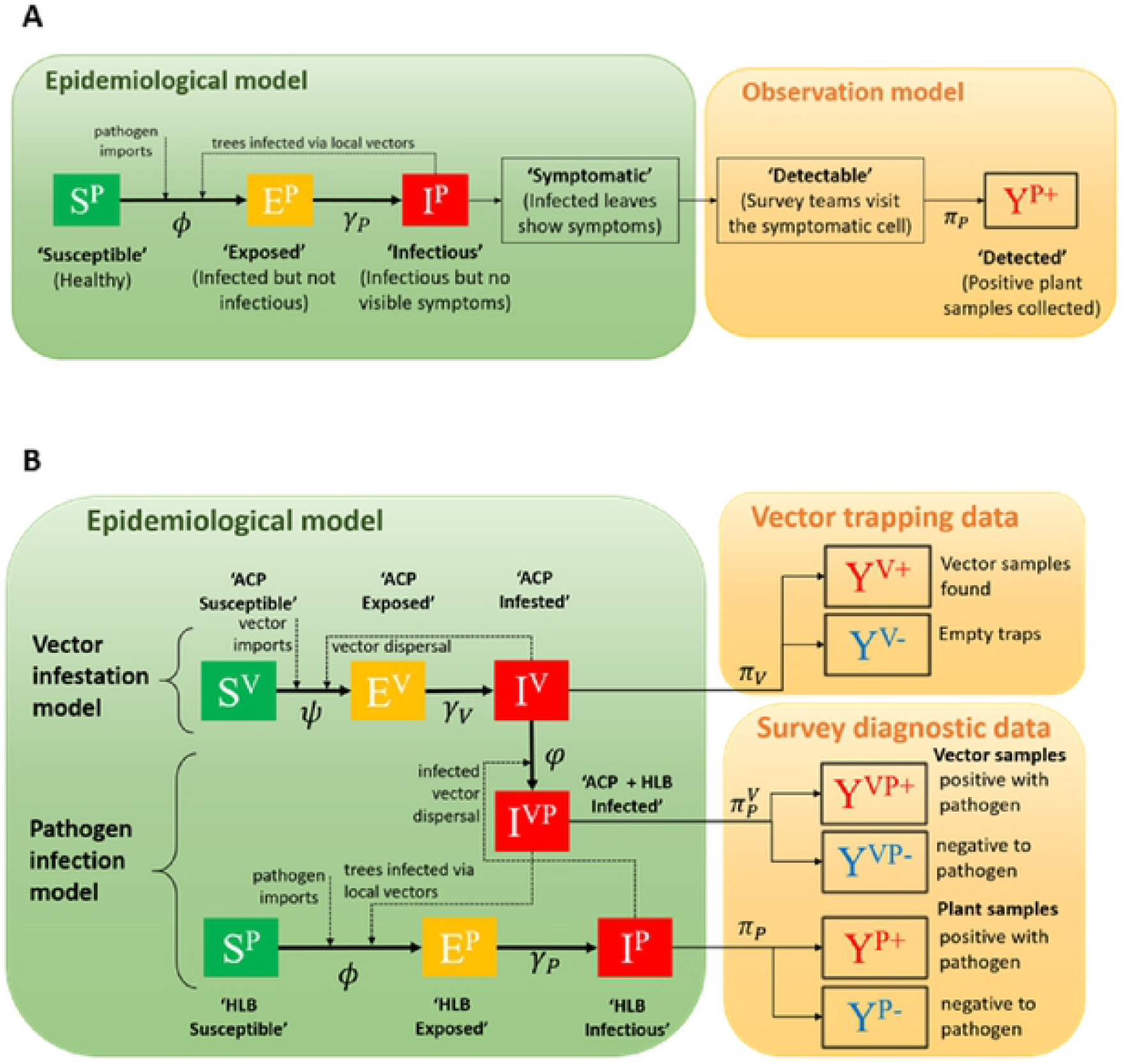
Epidemiological models for ACP and HLB spread. **(A)** The stochastic compartment model for HLB epidemiological dynamics in a Texas citrus grid cell and the observation model that matches infection status to survey diagnostic data. An infectious cell can infect other susceptible cells via the movement of local vectors, which were known to have established over the whole region before the emergence of HLB. **(B)** The joint epidemiological model for ACP and HLB dynamics in California where ACP is invading and not yet endemic. The model extends the Texas model and introduces a new epidemic category ‘ACP + HLB infected’ that connects the dynamics of ACP infestation to HLB infection in a grid cell.

Among the three sources of infection, the estimated rate of secondary transmission via local ACP vectors is three orders of magnitude greater than the rates of primary infection via cross-border infected vectors and four orders larger than human-mediated movement (Fig. 5A). The clear departure from zero of the posterior distribution for cross-border infection rate indicates the presence of surplus sources of infectious vectors over the Mexico border. Figure 5B presents the annual infection pressure caused by the three infection sources. We again observe the significant role of local vector movement in driving the epidemic, resulting in infection pressures an order of magnitude larger than primary forces in early years, rising to two orders in the later years. The posterior estimates for other parameters, including the dispersal scale, the average efficiency of the coordinated spraying program adopted by commercial growers and the waiting period from being infectious to detected for a citrus grid cell, are reported in Table 1 and Fig. S4A. We observed good agreement between the trained model’s simulations and the training and testing data for both temporal and spatial validation metrics (Figs. 4 and S4C). Besides, we also validated the model performance as a binary classifier (Fig. S4B) and evaluated the goodness-of-fit for various model variants (Figs. S2 and S3). Retrospective analyses of historical spread (Fig. 3) indicate a significant lag from the time that a cell became infectious until it is detected by the visual survey. Model simulations indicate that by October 2014, HLB had infected almost all of Hidalgo county (TX) whereas from the survey data it appeared as if the epidemic had just started to pick up. Having affirmed the model’s credibility and practicality for retrospective and future analyses for Texas, we adapt it to predict spread and management options for California.

**Fig. 3.**
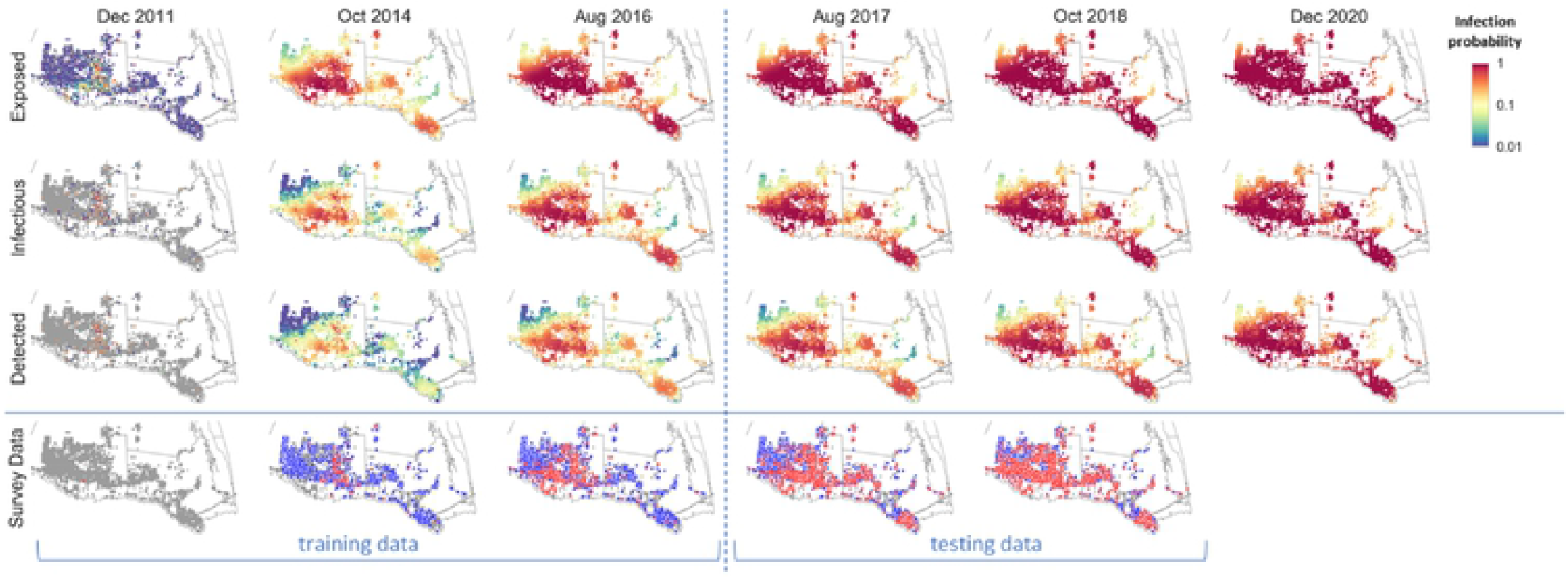
Spatiotemporal retrospective and prospective prediction of HLB spread in the Lower Rio Grande Valley, Texas. Spatiotemporal retrospective analysis of the historical spread of training period (Dec 2011 - Aug 2016) and prospective prediction of testing period (Sep 2016 - Oct 2018) and the future where no data were available (Nov 2018 - Dec 2020). We calculated infection risk by averaging over 1000 simulation realizations of the model fitted to the training data.

**Fig. 4.**
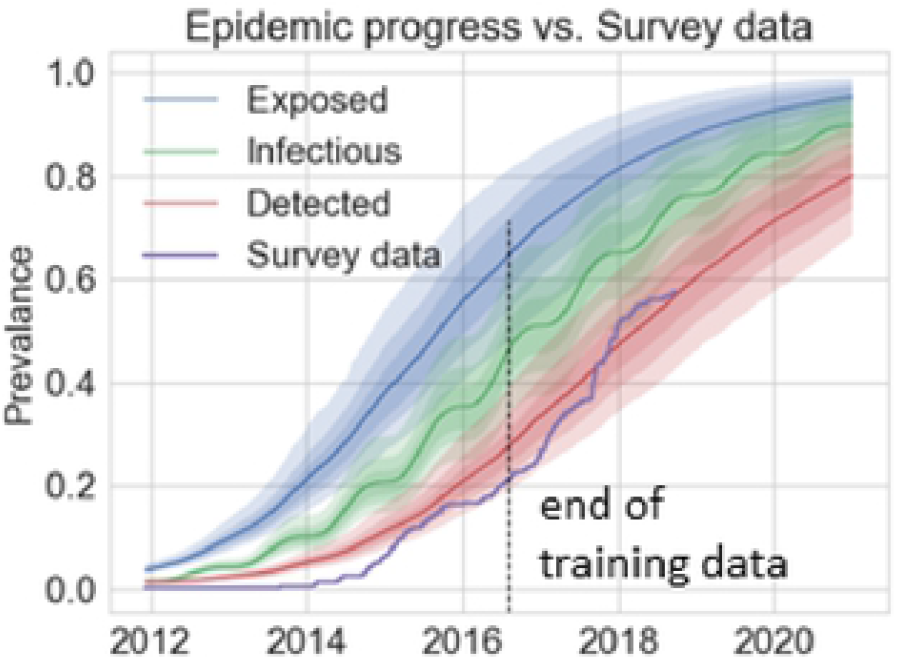
Temporal retrospective and prospective prediction of HLB spread in the Lower Rio Grande Valley, Texas. Temporal progression of the prevalence of three infection categories (Exposed, Infectious, and Detected) in comparison with the collected survey data. Besides the medians of 1000 simulation realizations (solid lines), we also show 50%, 75%, and 95% credible intervals (shades of decreasing intensities).

**Fig. 5.**
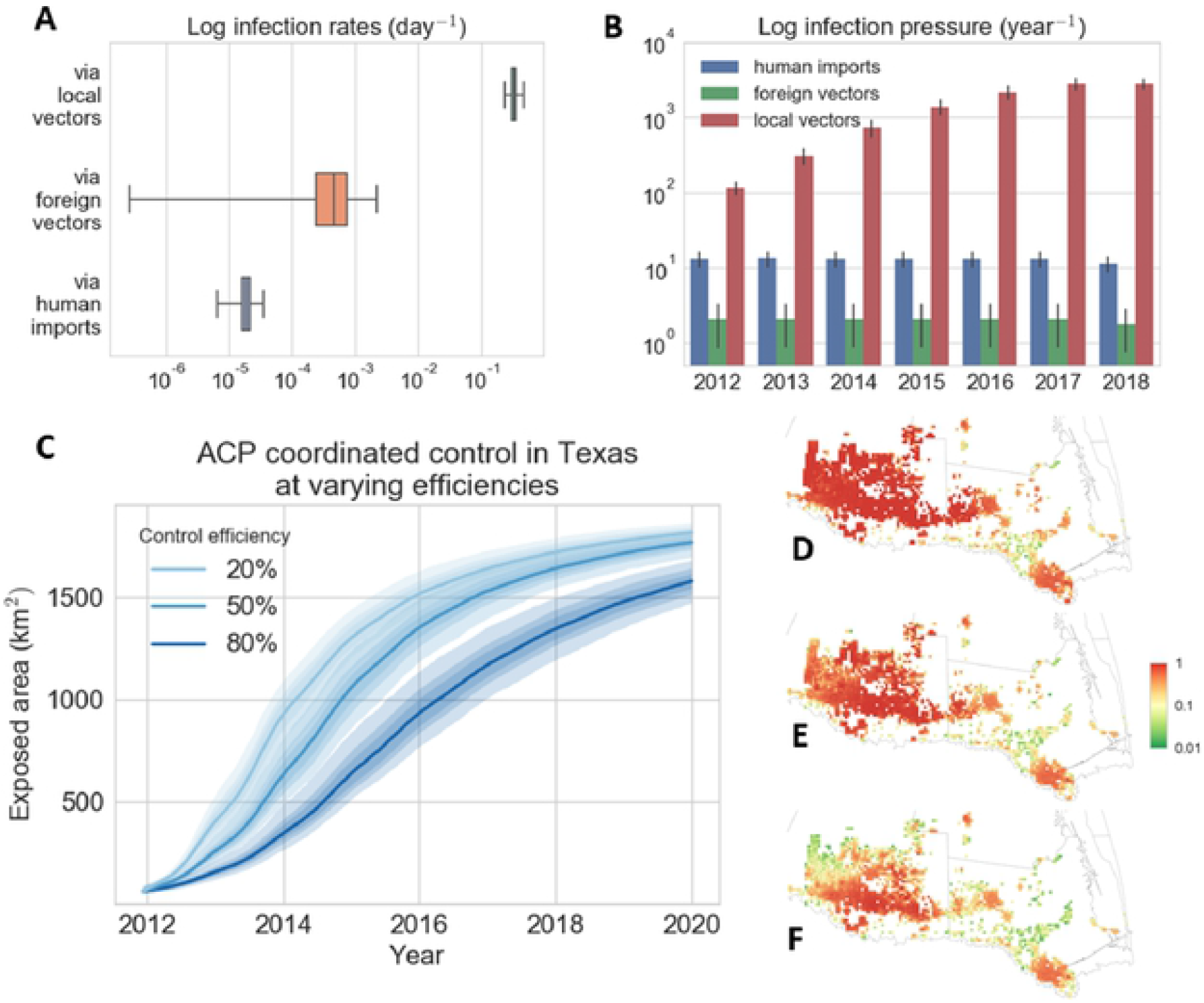
The role of primary and secondary forces of infection in Texas and the effectiveness of control scenarios in slowing down HLB spread. **(A)** Posterior distribution of infection rates via three sources: movement of local vectors from infectious trees in the landscape, the arrival of infected vectors over the international border, and introduction of infected trees by other human-mediated movement. **(B)** Contribution of the three transmission sources to the realized infection pressure overall on susceptible trees in the survey period 2012 - 2018. **(C-F)** The effect of the efficiency of the annual coordinated spraying program on (C) the temporal progression and (D-F) spatial snapshots in October 2014 of the cumulative Exposed area. We considered three different hypothetical area-wide coordinated control scenarios at efficiencies of (D) 20% (E) 50% and (F) 80%.

**Table 1.** Parameter description and their posterior estimates from Texas data. Unknown parameters for epidemiological and observation models and their posterior estimation. We used a Bayesian inference framework to probabilistically match model outputs with Texas HLB survey data and developed data-augmented MCMC algorithms to approximate parameter posterior distributions. Here we include the mean and 95% credible intervals of the acquired posterior distributions.

| Symbol | Description | Posterior estimates |
| --- | --- | --- |
| $\varepsilon_W$<br>(day<sup>-1</sup>) | Rate of HLB **primary infection** by external psyllids for cells far away from the Mexico border | $2.7_{0.1}^{14.2} \times 10^{-5}$ |
| $\varepsilon_B$<br>(day<sup>-1</sup>) | Rate of HLB **primary infection** by external psyllids for cells along the Mexico border | $4.65_{0.31}^{13.15} \times 10^{-4}$ |
| $\epsilon$<br>(day<sup>-1</sup>) | Rate of HLB **primary infection** by trade and other anthropomorphic activities | $1.90_{1.1}^{2.7} \times 10^{-5}$ |
| $\alpha$<br>(km) | Spatial **dispersal scale** for **ACP** movements | $1.96_{1.61}^{2.48}$ |
| $\beta_P$<br>(day<sup>-1</sup>) | Rate of HLB **secondary infection** | $0.33_{0.27}^{0.41}$ |
| $\beta_V$<br>(day<sup>-1</sup>) | Rate of ACP **secondary infestation** | $0.78_{0.61}^{1.03}$ |
| $\mu$ | Ratio of **feeding vs. moving** time of vectors | $5.14_{0.53}^{26.37} \times 10^{-3}$ |
| $\eta$<br>(%) | **Efficiency** of the annual coordinated **ACP spraying program** in commercial orchards | $79.6_{75.1}^{83.0}$ |
| $\sigma$<br>(day<sup>-1</sup>) | **Rate of HLB detection** by state-wide visual inspection survey | $2.13_{1.83}^{2.38} \times 10^{-3}$ |
| $\pi$<br>(%) | **Probability of detection of HLB** positive samples from an infectious site | $46.3_{44.6}^{48.1}$ |

### Prediction of emerging HLB spread in southern California

The HLB epidemic in southern California is at an earlier stage compared with the outbreak in Texas, with clusters of infected trees found in the Orange and Riverside Counties (Figs. 1B, 1E). Exploratory analysis showed that the data for Southern California are insufficient to provide a full set of parameter estimates for the landscape-scale model for California. The data are sufficient, however, to test the credibility of transferring Texas parameters to this new region. We accounted for the difference in weather conditions that affect ACP distributions in California and Texas by incorporating a weather, suitability score for ACP growth, particularly for temperature, (Liu & Tsai, 2000) to the model. We also introduced HLB quarantines and the removal of HLB-confirmed trees as discrete stochastic events into the spread model. We inferred the locations of unobserved infected cells up to 30^th^ June 2017 in southern California using the survey data collected before that date and simulated the epidemic forward. For the two years in the testing data (June 2017 - June 2019), the predicted detections successfully reproduced the spatiotemporal patterns observed in the survey data (Fig. 6). The predictions also showed good quantitative agreement with the testing data for both the temporal progression (Fig. 7A) and spatial autocorrelation metrics (Fig. S5A). Inspection of the model predictions for unobservable infectious categories (exposed and infectious cells) indicates that the extent of HLB spread is far greater than was detected by survey data. HLB is likely to be present in most counties in southern California and even with imposition of quarantines and tree removals upon detection, the disease is steadily increasing in severity. Predictions into the future suggest that by December 2021, HLB will have invaded the whole region of southern California. Our results indicate that surveys by visual inspection will reveal more HLB positive samples from all over Los Angeles and Orange Counties, and infected trees in San Bernardino, Riverside and San Diego Counties will become detectable.

**Fig. 6.**
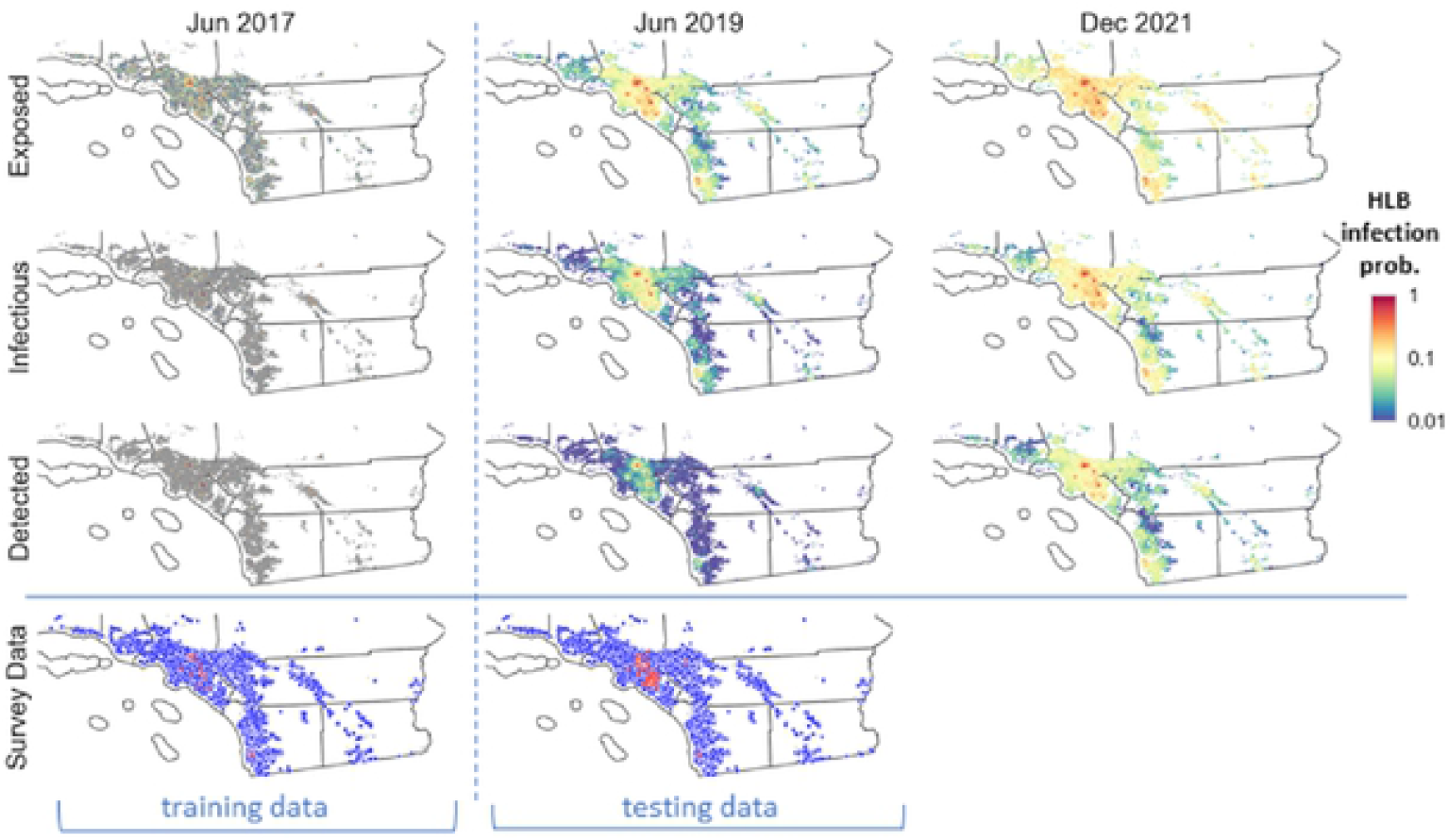
Spatiotemporal prediction of further HLB spread in southern California. We used California HLB survey data up to June 2017 to infer the locations of the hidden infectious cells and used the estimations to seed forward simulations. We calculated infection probabilities by averaging outputs from 1000 simulation runs for the prospective spread between June 2017 and December 2021.

**Fig. 7.**
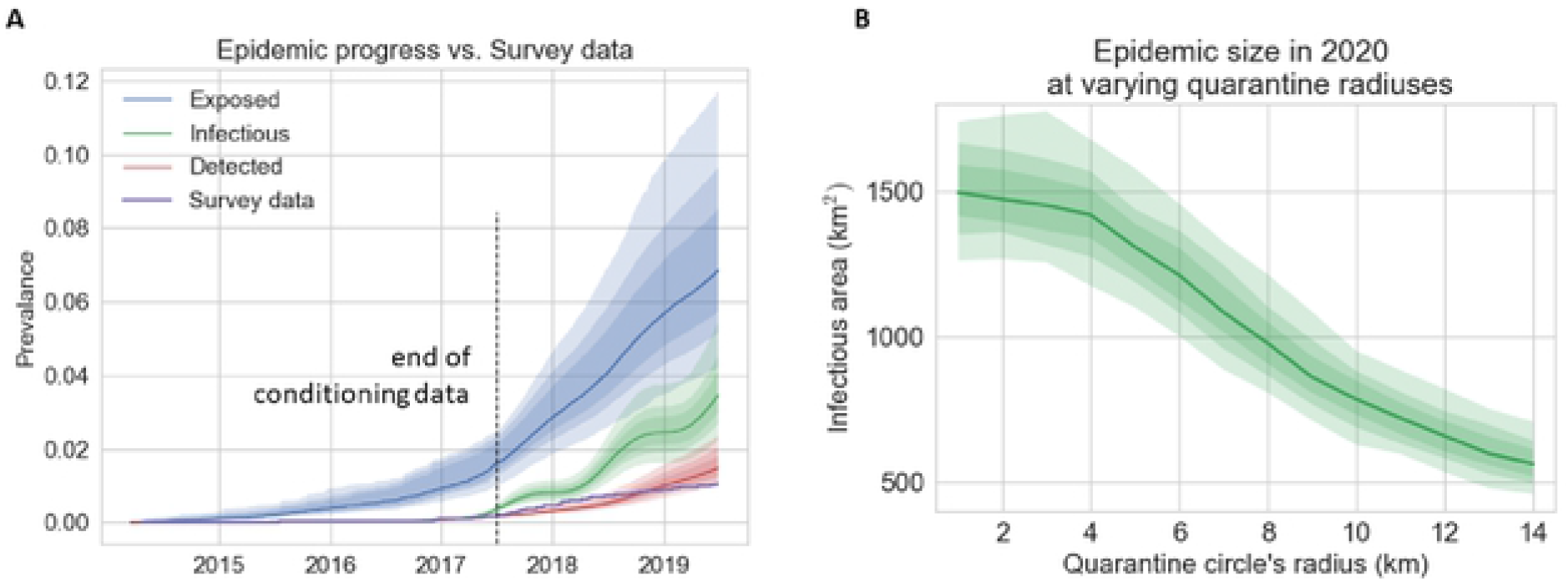
Evaluation of HLB epidemic progress in southern California vs survey data and the impact of quarantine radius to epidemic outcome. **(A)** Temporal progression of the prevalence of three infection categories (Exposed, Infectious, and Detected) in comparison with the training and testing data. We show means of 1000 simulation realizations as solid lines, and 50%, 75%, 95% credible intervals as shades of decreasing intensities. **(B)** The effect of the radius of the quarantine area centring around HLB newly-detected sites on the total Infectious area for southern California. We started simulations from June 2017 and assessed the total infectious area for December 2021.

### Bayesian inference of ACP invasion rate in the Central Valley using Texas survey data

The epidemic of HLB in the Central Valley, the major citrus growing area in California, is at an earlier stage than for southern California, with the vector, ACP, invading rather than being established (Figs. 1C, 1F). Modelling the spread for ACP at the landscape scale requires an estimate of the vector invasion rate. Exploratory analyses showed this was not possible using the small amount of ACP trapping data for the Central Valley: instead we calculated the ACP invasion rate using a more extensive ACP diagnostic dataset for Texas. Although ACP had fully invaded Texas by the start of data collection in 2011, the region was not fully infested with HLB-infected ACP. By introducing a new epidemic category ‘ACP + HLB infected’ that connects the dynamics of ACP infestation to HLB infection in a grid cell (Fig. 2B), it was possible to estimate an invasion rate parameter for ACP from the ACP diagnostic data collected as part of the Texas HLB survey (Fig. S1). In particular, the ‘ACP + HLB infected’ category marks cells containing HLB-infected ACPs. By modelling the transition of cells from ‘ACP infested’ to ‘ACP + HLB infected’, we estimated the rate at which vectors move from one cell to another.

We validated the ACP spread model by running simulations to reproduce the historic spread in the Central Valley in 2015 and 2016 and compared model predictions with the ACP trapping data, which were collected independently from the HLB survey and not used for parameter estimation. We incorporated the dynamics of the reactive ACP treatment program by the California Department of Food and Agriculture (CDFA) into the model and observed good agreement in both temporal progression (Fig. 9A) and spatial autocorrelation metrics (Fig. S5B), indicating that the ACP spread model successfully captures the ACP invasion dynamics.

### Prediction of potential HLB spread in the Central Valley as ACP is still invading

By conditioning the HLB spread model on the ACP spread model (Fig. 2B), we simulated joint spatio-temporal predictions of ACP and HLB potential spread in the Central Valley for the next ten years (Fig. 8). We hypothesise that regulators may consider dropping the reactive pesticide treatment by January 2025 as ACP will have invaded most residential areas in Fresno, Tulare, and Kern Counties and spread into the main commercial orchards in central Tulare County. The ACP and HLB epidemic will accordingly continue to expand even more rapidly (Fig. 9B). Our results indicate that by the end of 2030, ACP will have fully established over the whole Central Valley region, and a considerable number of HLB infected citrus trees will have presented in the main citrus growing areas (Fig. 8).

**Fig. 8.**
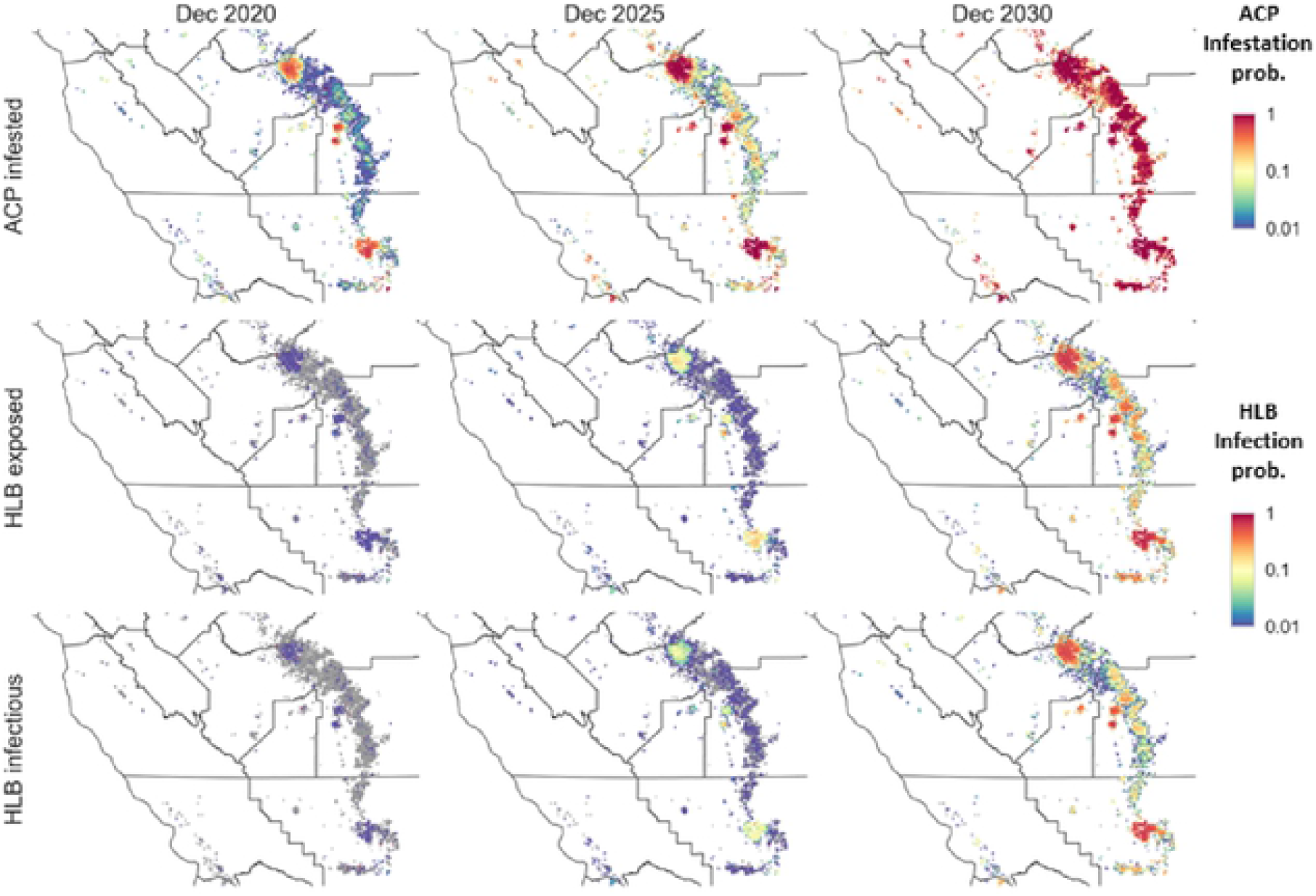
Spatiotemporal prediction of the potential ACP and HLB spread in the Central Valley. Weused Central Valley ACP trapping data to seed the simulations for ACP spread and locations of inconclusive HLB samples (Ct value less than 38 in qPCR diagnostic test) as the initial infected HLB sites. HLB spread can only happen between ACP infested sites. We calculated the ACP infestation and HLB infection probabilities by averaging over 1000 simulation runs for the prospective epidemics from January 2020 to December 2030.

**Fig. 9.**
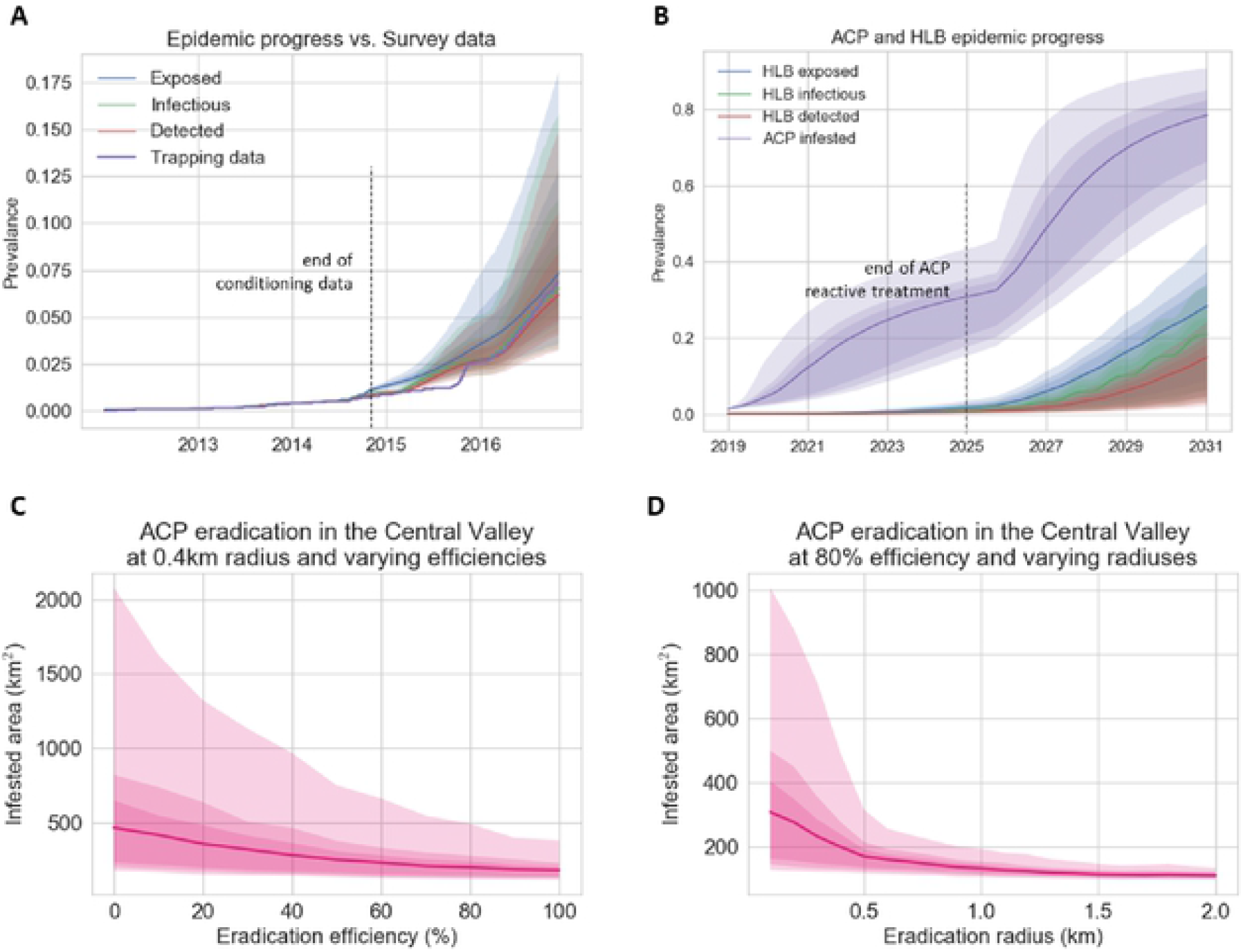
Potential ACP and HLB epidemic progress in the Central Valley and the likely effectiveness of control. **(A)** Temporal progression of the predicted infestation prevalence in comparison with the trapping data for the data availability period (up to October 2017). **(B)** Temporal progression of the predicted ACP and HLB epidemics from January 2019 to December 2030. We assumed that as ACP prevalence passes 0.3, ACP the reactive treatment program would have been dropped due to the high cost of maintaining the program and the reduced effectiveness as ACP becomes widespread. **(C, D)** The effect of varying the efficiency and radius of pesticide treatment upon detection of the vector ACP on the total Infested area in the Central Valley. We started simulations from January 2020 and calculated the total Infested area for December 2021 for (C) control efficiency from 0% to 100% for treated circles of radius 0.4 km and (D) treatment radius from 0.1 km to 2 km assuming spraying efficiency of 80%.

### Evaluation of the impact of putative control scenarios in Texas and California

Besides providing predictions of vector and pathogen spread, the models can be used to evaluate the impact of putative control strategies for containing or slowing down the HLB epidemic. We ran retrospective and prospective simulations of the ACP and HLB spread models to assess the epidemic impact of the control scenarios most relevant to each region: the coordinated ACP spraying program in Texas (Figs. 5C-F), the HLB quarantines program in southern California (Fig. 7B), and the reactive ACP treatment in Central Valley (Figs. 9C, D).

The area-wide coordinated vector spraying program in Texas was carried out annually since the start of the HLB epidemic in 2011. Growers joined the program by applying pesticide sprays within a short, designated period to target the overwintering vector populations. Using historic HLB survey data, we estimated that the program helped to reduce about 80% (designated as the control efficiency) of the vector population in commercial orchards. To understand the importance of having an area-wide collaborative effort amongst growers in place, we ran retrospective simulations for hypothetical scenarios in which less intensive ACP control had been carried out (Figs. 5C-F). We observed a nonlinear relationship between control efficiency and HLB exposed area. While increasing control efficiency from 20% to 50% cannot guarantee the reduction of epidemic size, bringing control efficiency up to 80% was effective (we observed both a significant reduction of the infected area in 2020 and a clear separation of the 95% credible intervals associated with 20% and 80% control efficiency). Having established the importance of maintaining a coordinated spraying program, we considered the impact of increasing the size of HLB quarantine areas surrounding confirmed positive sites in southern California (Fig. 7B). Current regulation imposes a 5-mile (8 km) quarantine radius around each HLB positive site and restricts the movement of citrus tree products from the quarantined area to other places. Simulation results demonstrated consistent reduction of the infectious regions as the quarantine radius is increased, with the currently prescribed 8 km radius helping to reduce a third of the epidemic size at the end of 2021.

As the current focus for the Central Valley is on ACP invasion, we considered two parameters that drive a reactive ACP eradication program: the eradication efficiency (Fig. 9C), and the radius of the treated circle around an ACP positive site (Fig. 9D). Our simulations allowed for pesticide treatment, applied by CDFA, on all citrus trees within a 400 m radius of an ACP positive site. Where treatment circles overlap a commercial grove, the whole grove is treated. Reactive treatment occurs in addition to a coordinated spray implemented annually by commercial growers. Simulation results show that having both high efficiency and sufficient radius are essential to slow down the spread of ACP in the Central Valley (Figs. 9C, D). There is a consistent reduction in the median infested area and also the 95% credible interval as the eradication efficiency increases. An eradication efficiency of 80% reduces the median infested area by half. It also reduces the upper boundary of the credible interval by 75% compared with no treatment. Increasing the eradication radius from 100 m to 500 m decrease the expected infested area by 50% and the credible interval by 75%. The results suggest that although 80% eradication efficiency is reasonable, it might be worthwhile to increase the treatment radius to 500 m.

## DISCUSSION

We have developed, parameterised and tested a unified and flexible epidemiological modelling framework to predict the spread of the Huanglongbing (HLB) disease on citrus at landscape scales. Our stochastic framework takes account of the intrinsic uncertainty in disease spread. The framework allowed us to construct predictive models using comparatively sparse data from early surveys as the disease spreads into a new region. We used it to screen effective management strategies at landscape scales, advancing previous models that focused on individual trees (Lee, et al., 2015) and plantations (Parry, et al., 2014).

We use conventional compartmental models to account for unobserved latent and cryptic infection as well as symptomatic infection. The framework allows us to accommodate the impact of disease control measures on the surveillance data used when estimating transmission and dispersal parameters for the pathogen. The framework also accommodates two scenarios, one in which the insect vector (the Asian citrus psyllid: ACP) is endemic, the other when the vector is also invading a new region. The main practical outcome of the work is to predict the spread and potential for control of HLB in southern California, where the pathogen has been reported and the insect vector is endemic, and in the major citrus production region in the Central Valley in California, where the vector is spreading but the pathogen has not yet been reported.

Building early predictive models is critical to disease management for outbreaks of infectious diseases. For example, thanks to the early availability of case data for the 2001 foot-and-mouth disease (FMD) outbreak in the United Kingdom, researchers were able to develop mathematical models quickly and used them to formulate control policies (Keeling, et al., 2001; Ferguson, et al., 2001). Similar extensive control strategies had been examined for the severe acute respiratory syndrome (SARS) epidemic in Hong Kong (Riley, et al., 2003). The level of reporting for human and animal diseases is generally higher than for plant epidemics. The situations in California exemplify a common problem in plant epidemiology, where data are scarce (McRoberts, et al., 2019) and there are too few observations on disease and vector spread to infer parameters from which to predict spread within the locations at risk. Here, we use well-documented surveillance data from Texas for parameter estimation, model selection and validation. The data were collected from the onset of the epidemic for commercial and residential areas in south Texas until HLB had widely invaded. The distribution of infected vectors was also monitored. The Texas data were preferred to surveillance data from Florida, where the disease is now widespread, because the data reflect times at which locations were sampled rather than the intrinsic spread of HLB in that state (T. R. Gottwald, pers. comm.). We incorporate differences in daily temperatures and the effect of ongoing control strategies in transferring the model from Texas to the Californian situations in model simulations.

It is challenging to estimate epidemiological parameters from surveillance data on populations that are exposed to disease control. In the case of Texas, coordinated spraying was applied amongst commercial orchards to kill the insect vector. We used a Markov chain Monte Carlo method with data augmentation (DA-MCMC). The DA-MCMC method is considered a robust approach for inferring parameters for stochastic, individual-based epidemiological models (Gibson & Renshaw, 1998; Jewell, et al., 2009; Parry, et al., 2014; Keeling, et al., 2021). The method has been used to estimate epidemiological parameters for heterogeneous large-scale epidemic systems with cryptic infections. Applications include foot-and-mouth outbreaks on cattle (Jewell, et al., 2009), avian influenza epidemics on poultry (Jewell, et al., 2009), and MRSA outbreaks in hospital wards (Kypraios, et al., 2010). Convergence of the DA-MCMC method, however, is known to be difficult when applied to domains (landscapes) with heterogeneously distributed target populations. Accordingly, we improved the mixing and convergence of MCMC samplers for unobserved epidemic transitions by utilising the randomised construction of Markov trajectories (Gross & Miller, 1984) and exact inference algorithms for hidden Markov models (Rabiner & Juang, 1986). The improved algorithms allowed us to successfully reproduce the HLB epidemiological dynamics and control interventions observed in Texas, and retrospectively to analyse the impact of varying efficiency of the coordinated ACP spraying program on HLB epidemics.

Using the improved MCMC samplers, it is possible to infer locations that are cryptically (i.e. asymptomatically) infected from the survey data available at the time of prediction. Initialising spatiotemporal epidemic models with asymptomatic as well as symptomatic infected sites is essential to capture the current extent and the future potential for disease spread. We used the Texas HLB survey data up to 2016 as a training dataset for parameter estimation and the remaining data up to 2018 for model validation. When tested on data not used in parameter estimation, the HLB spread model showed remarkable agreement for both spatiotemporal visualisation and temporal and spatial evaluation metrics (Fig. 3). Retrospective inference of HLB infection times for Texas showed that the epidemic progressed at a much faster pace than had been captured by the survey data. Knowledge of the locations of cryptically infected sites gives government and industry decision-makers a two-year advantage in knowledge of the extent of the epidemic when compared with survey data alone. We were able to infer the average efficiency of the annual coordinated spraying from the Texas survey data and to run retrospective simulations to assess the impact of having such measures in place in comparison with less effective executions of the program. Our results showed the benefit of having a high level of participation from commercial growers in slowing epidemic progression, especially during the first few years after the invasion (Fig. 5C).

Allowing for cryptically infected sites (estimated from the training data) as well as survey reports of symptomatic sites when initialising the HLB spread model in southern California gave very good agreement in predicting the spread patterns observed in the two years of test data (Fig. 6). The models were used to investigate the impact of different control strategies on epidemic outcomes for HLB in California. Increasing the radius of the quarantine area to prevent movement of citrus products around newly-detected HLB sites has the potential to reduce the total Infectious area for southern California (Fig. 7B). Within the Central Valley, our results indicate that changing the radius and the efficiency for the reactive ACP treatment programme each reduce the infested area (Fig. 9C, D). There was also a marked reduction in the uncertainty of the outcomes of the programmes with enhanced control effort (Fig. 9C, D).

The modelling and inference framework described in this paper can easily be adapted and extended to tackle various epidemiological problems. An obvious next step is to incorporate the operating costs and potential economic gain for the putative control strategies considered for southern California and the Central Valley. Predictive analysis using model simulations similar to those used in the paper can provide recommendations for optimal control strategies by decision-makers. An important aspect of the voluntary coordinated ACP spraying program is to evaluate the impact of grower behaviours in terms of compliance and responsiveness to the efficiency of reducing the spreading power of ACP populations. Adapting the epidemiological models for a retrospective analysis of past HLB epidemics, e.g., in Florida, should allow for valuable insights about what could have been done to slow down the epidemic course. As the African citrus psyllid, *Trioza erytreae* (Del Guercio), in Portugal and Spain constitutes a threat of introduction of the disease into European countries, the framework can be used to transfer U.S. parameters to these countries pending enough data for local estimation, by accounting for differences in psyllid behaviours, weather suitability, as well as regional operations.

## METHODS

Here we provide a summary of the data sources, epidemiological models, parameter estimation methods, and prediction procedures used in the paper. Please consult the supplementary information for more detailed descriptions.

### Data sources

We obtained data from surveys for early HLB detection in Texas and California and detection of ACP presence in California. The data were collected and analysed by the U.S Department of Agriculture (USDA) and the California Department of Food and Agriculture (CDFA). Data collection surveys were part of an intensive area-wide management research program for citrus greening over multiple U.S. states and localized management of ACP in California. We used the HLB survey data in Texas (Fig. 1A, D) to estimate parameters and validate the HLB epidemiological model (Fig. 2A) We also used an additional survey of trapping data for HLB-infected ACP in Texas to estimate parameters for vector spread (Fig. S1B in Supplementary Information). HLB survey and ACP trapping data for southern California (Fig. 1E) and Central Valley (Fig. 1F) were used to initiate simulated epidemics for model inference on spread and control strategies and further validation. All data were geo-mapped and aggregated by visiting dates to obtain spatiotemporal datasets of HLB positivity for plant and vector samples in Texas (Figs. 1D and S1B), plant samples in southern California (Fig. 1E) and ACP presence in the Central Valley (Fig. 1F).

### Epidemiological models

We developed a spatially explicit, continuous-time, stochastic compartment model for the HLB spread in Texas and southern California under the assumption that the vector had already become endemic. The landscapes were rasterised into 1 km x 1 km grid cells. Each grid cell occupied by citrus trees belongs at any one time to one of the four infection categories: HLB susceptible, exposed, infectious, and detected (Fig. 2A). We extended the model to account for the fact that the underlying ACP population in the Central Valley is still spreading and has not fully invaded the region (Fig. 2B). In addition to the above four HLB infection compartments, we consider three compartments for ACP infestation status in grid cells: ACP susceptible, exposed and infested. We also introduce a new epidemic category to associate the dynamics of ACP infestation to that of HLB infection: ACP + HLB infected. This category marks cells that observe the presence of HLB-infected vectors.

A susceptible cell is exposed to HLB infection when the first tree in the cell is infected via three transmission sources. Primary transmission can arise from the introduction of infected trees and tree products by trade and human-mediated movements (represented as parameter *ϵ*) or from HLB-carrying vectors arriving from external environments (represented by two parameters away from, *ε_W_*, and across the US-Mexican border, *ε_B_*). Using two different parameters to represent the second primary transmission source reflects our assumption that citrus trees along the Mexico border were under extra epidemic pressure from a higher incidence of infected insects on the Mexican side of the border compared with those further from the border. Secondary transmission from an HLB infectious to a susceptible cell happens by the flux of vectors moving between the two cells. We denoted *β_p_* as the rate for secondary infection and used an isotropic exponential kernel, *K_α_*(*r*) ∝ *e^-r/α^* where *α* represents the dispersal scale, to depict the dependence of movement rate on the spatial distance *r* between the cells. We derived the HLB exposure rate, given in equation (1) in the SI, using a mechanistic model for vector movement and feeding. Besides the variation in citrus density over the landscape, the model also accounts for the variation in vector density that is affected by the annual coordinated spraying program to commercial citruses and daily weather. We used a parameter *η* to represent the efficiency of vector control and the weather-driven model for vector development rate by (Liu & Tsai, 2000). The parameters *ϵ, ε_B_, ε_w_, α, β_P_, η* were estimated using the Texas HLB survey plant diagnostic data.

In the Central Valley, a susceptible cell is exposed to ACP infestation when the first few vectors arrive in the cell from nearby sites (represented as secondary infestation rate, *β_v_*) or transported from external environments (represented as primary infestation rate *ε_v_*). The force of ACP exposure was derived using the same vector flux model as in the case of HLB exposure rate and is given in equation (2) in the SI. The challenge, however, lies in estimating the secondary infestation rate *β_v_* as we cannot infer it directly from California ACP trapping data. We observed that the rate at which infected vectors migrate from an ‘ACP + HLB Infected’ cell to an ‘ACP Infested’ site in the extended model (Fig. 2B) is equivalent to the secondary infestation rate *β_V_* under the assumption that Las-carrying vectors behave similarly to uninfected vectors. Therefore, we can derive the transition force (equation (3) in the SI) and estimate the parameter of interest using the vector diagnostic data from the Texas HLB survey. Since we carried out parameter estimation under a Bayesian inference framework, we can use the previously acquired posterior estimates for *α,μ,η* to complement the sparsity of vector diagnostic data.

We prescribed a latent period for a cell to transition from exposed to infectious to HLB and from exposure to infested to ACP. We used a seasonally-forced model for the rate of infectiousness/infestation onset (Parry, et al., 2014). Parameters for the latent periods were obtained from in-orchard and in-nursery observations.

### Parameter estimation

We adopt a Bayesian approach to estimate parameter values from noisy survey data. We treated the unknown timing of epidemiological transitions as random variables and used a data-augmented MCMC algorithm to infer about them. An MCMC algorithm approximates the joint distribution of parameters by constructing a Markov chain that converges to the desired distribution in equilibrium. After a burn-in period, the samples generated from the chain form an unbiased representation of the parameter posterior distribution. We used the Metropolis-Hasting method (Chib & Greenberg, 1995) to construct samplers for both parameters and unobserved epidemic transitions. We used the randomized construction of Markov trajectory (Gross & Miller, 1984) and exact inference algorithms for hidden Markov models (Rabiner & Juang, 1986) to improve samplers of epidemic transitions. For more details of the algorithms, please consult the SI.

The overall likelihood of a set of parameter values comprises the model likelihood and the data likelihood. The model likelihoods (equations (5) and (8) in the SI) can be naturally derived from the stochastic construction of the HLB and ACP epidemiological models described above. We developed data likelihoods (equations (6, 7, 9) in the SI) using two parameters of the data collection process: *π* represents the probability that a positive sample is collected from an infectious cell, and *σ* indicates the expected duration from becoming infectious to getting detected. Both parameters, together with parameters of epidemiological models, were estimated from Texas HLB survey data.

### Model validation and prospective prediction

We estimated parameters using Texas HLB survey data collected between December 2011 and August 2016. We then validated region-specific epidemiological models using either a different survey time range or an independent source from the training data. In particular, we used data collected between September 2016 and October 2018 as the testing data for Texas and data collected in southern California from June 2015 to June 2019 to test the region HLB model. ACP trapping data in 2015 and 2016 were used to verify that the ACP spread model for the Central Valley agrees with reality. We compared model simulations to testing data in terms of both temporal progression and spatial autocorrelation metrics. Please consult the SI for more details of validation construction.

To analyse the sensitivity of different model components to prediction performance, we considered four variants to the full model described above. Each model variant differs from the full model by one component of the secondary transmission model: (1) no normalisation, in which the normalisation term for vector fluxes is assumed to be the same for all cells and absorbed to the secondary infection rate; (2) no control effect, in which we ignored the occurrence of the annual coordinated spraying program; (3) no border effect, in which we did not distinguish between sites near to and far from the Mexico border and used the same primary infection rate for infected vector from external environments; (4) power-law kernel, in which the exponential dispersal function is replaced with a power-law function. Model variants were fitted using Texas HLB survey data up to August 2016 and validated with data up to August 2017 (Figs. S2 and S3).

We made prospective predictions more than two years past the final observation time for each region. Not all sites already infected with HLB are observable at the time of forecast. We used observed survey data to infer the locations of cells that have been exposed to and infectious with HLB at that time. We used an MCMC-based simulator analogous to the data-augmented MCMC algorithms used for parameter estimation to sample epidemic transition times that agree well with observed data. The effect of known control measurements carried out in each region was incorporated using various mechanisms (details in the SI). These include the annual coordinated spraying program in Texas, reactive removal of infected trees and HLB quarantine in southern California, and reactive vector spraying upon ACP detection in sticky traps in the Central Valley. Besides forecasting into the future, the prediction models can also be used to evaluate the impact of putative control strategies by varying parameter values that drive control measurements.

## Acknowledgements

This material was made possible, in part, by a Cooperative Agreement from the United States Department of Agriculture’s Animal and Plant Health Inspection Service (APHIS). It may not necessarily express APHIS’ views.

We would like to thank Tim Gottwald, Nik Cunniffe, Sunil Kumar and members of the Epidemiological and Modelling group for useful comments and discussions. CAG acknowledges support from the Bill and Melinda Gates Foundation.

